# Selective Activation of Nerve Fiber Subpopulations with Intrafascicular Stimulation

**DOI:** 10.64898/2026.05.27.727943

**Authors:** Arianna Ortega Sanabria, Louis Regnacq, Anil K Thota, Avery Holmes, Justin M Asbee, Sylvie Renauld, Florian Klobl, Yannick Bornat, Samantha Robinson, Laura M McPherson, James J Abbas, Ranu Jung

## Abstract

**Background:** Peripheral nerve stimulation (PNS) is most effective when specific nerve fiber subpopulations are activated, while minimizing off-target activation, which may cause undesirable side effects. This selectivity depends primarily on electrode design and charge delivery. We hypothesized that selective PNS could be achieved through electrode placement and intrafascicular electric field steering using Longitudinal Intrafascicular Electrodes (LIFEs).

**Methods:** LIFEs were implanted into the tibial fascicle of the sciatic nerve of 17 anesthetized adult rats. We tested whether electrodes positioned at different cross-sectional and longitudinal locations within the same fascicle, together with different electric field-steering approaches produced distinct activation patterns in the gastrocnemius lateralis muscle. Muscle responses were measured using high-density epimysial electromyography (HD-eEMG).

**Results:** Electrodes placed at different locations within the same fascicle activated distinct muscle regions, demonstrating intrafascicular selectivity. Bipolar stimulation recruited nerve fibers differently than monopolar stimulation, showing that electric field steering can further shape the selective recruitment. In both configurations, increasing the stimulation amplitude produced a graded increase in muscle activation. Furthermore, our findings demonstrated that HD-eEMG is an effective tool for evaluating intrafascicular selectivity.

**Conclusion:** These findings suggest that improving on-target selectivity may support next-generation bioelectronic therapies with better outcomes and fewer side effects, potentially enabling more precise, organ-specific neuromodulation. Using multiple intrafascicular electrodes may provide two complementary strategies for enhancing selectivity: strategic intrafascicular placement to access different fiber subpopulations and bipolar configurations to steer recruitment beyond what a single electrode can achieve.

## 1. Introduction

Peripheral nerve stimulation (PNS) is a promising therapy for conditions inadequately managed by pharmaceuticals or physical rehabilitation, including depression, epilepsy, lower back pain, and bladder or fecal incontinence (1–4). However, its clinical utility is limited by its off-target effects. For example, vagus nerve stimulation (VNS) can treat depression (1,5) and reduce epileptic episodes (3); however, side effects, such as hoarseness and neck pain often lead to discontinuation (4). Similarly, sacral nerve stimulation (SNS) in inflammatory bowel disease and urinary or fecal incontinence (6) can cause abnormal sensations, abdominal pain, altered intestinal function, and pelvic tingling (2). In individuals with upper-limb amputations, cuff-based PNS can evoke useful sensory percepts but may also trigger unwanted motor activity (7).

A major cause of these off-target effects is limited stimulation selectivity, which leads to the unintended activation of non-targeted nerve fibers. This limitation is partly due to an incomplete understanding of nerve structure and organization (8–10). Improving selectivity is therefore essential to reduce side effects, improve outcomes, and enable more precise, organ-specific neuromodulation therapies (8,11,12).

Neurostimulation selectivity can be categorized into two primary types: fiber-type and topological selectivity. Fiber-type selectivity refers to the preferential activation of specific classes of nerve fibers, such as motor or sensory fibers. In contrast, topological selectivity involves spatially targeted stimulation, either at the level of fascicles (interfascicular selectivity) or within subpopulations of fibers in a single fascicle (intrafascicular selectivity) (8,13). This study focuses on intrafascicular topological selectivity.

Selectivity also depends on the neural interface, including intraneural, extraneural, or surface electrodes (8,14,15). Although extraneural cuff electrodes are widely used because they are easy to implant and are clinically effective, their distance from the target fibers limits their selectivity. In contrast, positioning electrodes closer to the target population improves selectivity (8,11,16). Consequently, intraneural electrodes, such as longitudinal intrafascicular electrodes (LIFEs), are particularly promising for enhancing intrafascicular selectivity because they are placed within the fascicle near specific fiber groups (11,17,18).

Another approach to improving interfascicular selectivity is extraneural field steering, which uses multiple electrode configurations to shape the electrical field more precisely (19,20). This approach can selectively activate different fascicles and improve interfascicular selectivity (20–23). However, targeting centrally located fascicles remains limited and has not been shown to substantially improve intrafascicular selectivity (24).

In this study, we investigated the effects of electrode placement and intrafascicular field steering on nerve fiber recruitment using single and multiple LIFEs implanted within the tibial fascicle of the sciatic nerve. High-density epymisial electromyography (HD-eEMG) arrays were used to assess the resulting muscle activation patterns. Our results demonstrate that deploying multiple LIFEs within the same fascicle in a monopolar configuration enables selective activation of distinct subpopulations of nerve fibers, resulting in different spatial muscle activation patterns. Furthermore, using multiple LIFEs in a bipolar configuration enables field steering, shifting the focus of activation, and altering the spatial muscle activation pattern.

These findings indicate that electrode placement within the fascicle and field steering can enhance intrafascicular selectivity. This approach has the potential to advance focal neural stimulation and support the development of organ-specific neuromodulation therapies, with minimal or no side effects.

## 2. Methods

We investigated two strategies for enhancing intrafascicular stimulation selectivity using LIFEs. Specifically, we assessed the effect of positioning multiple LIFEs within the same fascicle and using multiple LIFEs within the same fascicle arranged in a bipolar configuration to steer the electric field. The experiments were performed on 17 adult male Sprague-Dawley rats (433 ± 29 g) under anesthesia. All animal care and handling procedures were conducted in accordance with protocols approved by the Institutional Animal Care and Use Committee of the University of Arkansas (Protocol AUP22014).

### 2.1 Electrode Fabrication

Custom LIFEs were fabricated from insulated Pt/Ir wires (27.5µm in diameter) with an active site of approximately 1mm created by de-insulating the wire. One end of the wire was welded to a tungsten needle (75µm in diameter) to facilitate fascicular implantation, whereas the opposite end was welded to a metal plate to connect to the stimulator. The electrode impedance was verified using a custom bioimpedance measurement system (25), ensuring values within 8 – 40 kΩ (mean: 16.9 ± 9.5 kΩ) at 1kHz. From these recordings, we estimated the active site length of the electrodes to be 550 ± 100µm (26).

### 2.2 Surgical procedures and electrode implantation

Anesthesia was induced and maintained with inhaled isoflurane (1-3% ISO in O_2_, flow rate approximately 3L/min). The sciatic nerve was exposed via a lateral thigh incision, and the surrounding muscles were gently retracted using elastic stays to isolate the nerve.

Figure 1 illustrates the design of the *in vivo* experimental study. LIFEs were implanted into the tibial fascicle of the sciatic nerve (2-4 electrodes per rat; Figure 1a), aligned parallel to the axons, by inserting the needle from the proximal to distal end using a custom implantation tool. Appropriate placement and electrode functionality were verified by eliciting paw movements using a custom-built stimulator (27) that delivered biphasic, symmetrical square pulses. Electrodes that failed to evoke paw movement were excluded from stimulation. Electrodes were secured to the epineurium using 8-0 non-absorbable sutures (DemeTECH®). The paw was stabilized using a 3D-printed fixture.

**Figure 1.**
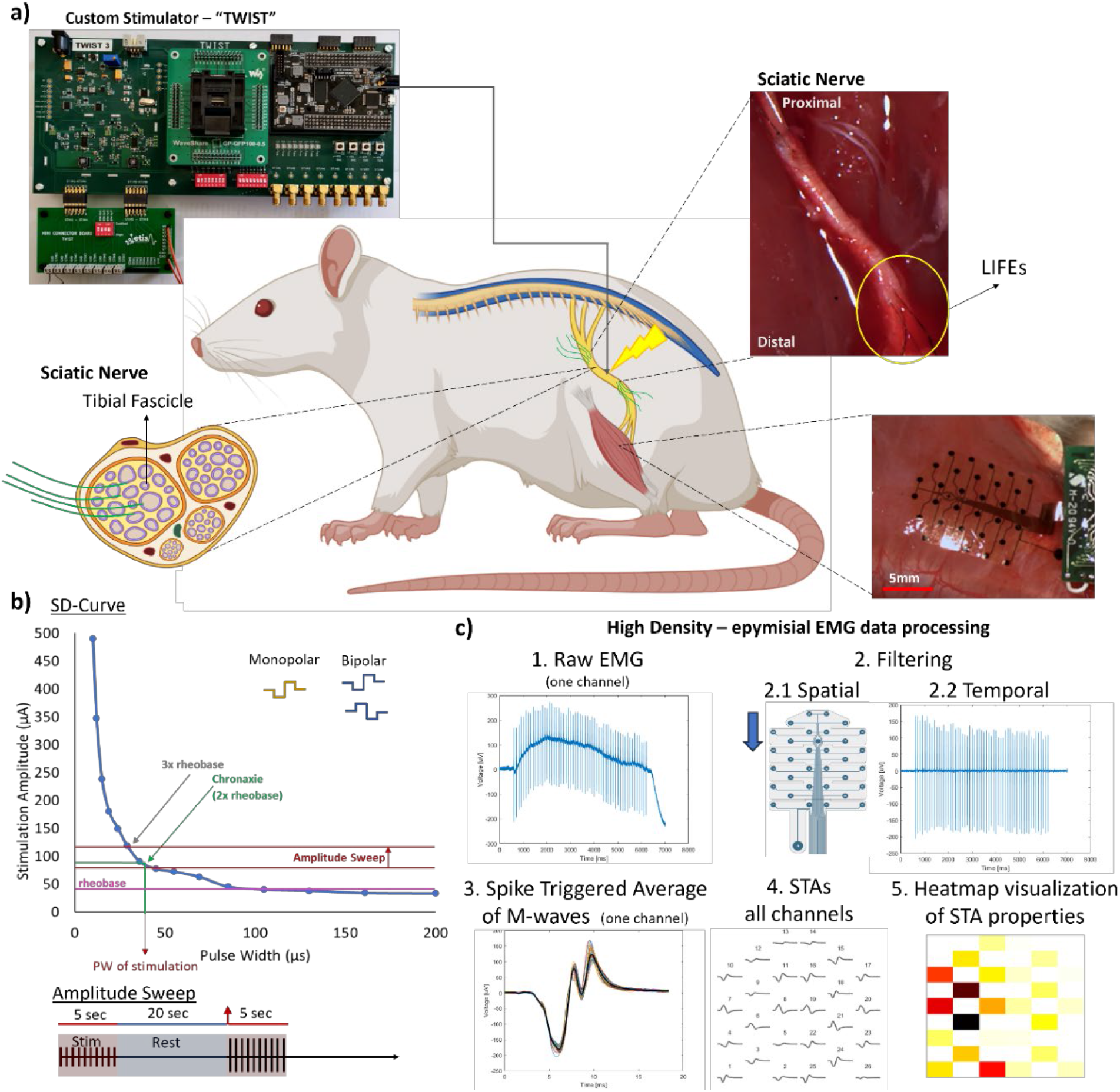
Study design. (a) Schematic: A custom stimulator “TWIST” generated electrical stimulation pulses delivered to the tibial fascicle of the sciatic nerve using longitudinal intrafascicular electrodes (LIFEs). High-Density Epimysial electromyograms (HD-eEMG) are collected from the gastrocnemius lateralis muscle with a flexible array. Schematic partially made with Biorender.com (b) Strength-Duration (SD)-Curves were used to identify the rheobase and chronaxie for each electrode and stimulation configuration. To generate SD-Curves, pulses at 1Hz were delivered at each pulse width and the stimulation amplitude was increased until a muscle twitch was visible. The amplitude sweep protocol consisted of stimulation pulses delivered in monopolar or bipolar configurations in bursts of 5 seconds interspersed with 20 seconds rest to prevent muscle fatigue, with increase in pulse amplitude in subsequent bursts. (c) The HD-eEMG raw signals were spatially and temporally filtered. Subsequently, stimulus-triggered averages (STA) were calculated from the M-wave responses for each HD-eEMG channel. The root mean square (RMS) amplitude derived from the STA was used to generate heatmaps, providing a visual representation of the spatial distribution of muscle activation.

Body temperature was continuously monitored using a rectal thermometer, maintained throughout the experiment with a heating pad (Kent Scientific, Inc.), and supplemented with hand warmers as needed. Heart rate, respiratory rate, and SpO_2_ were continuously monitored (Vetcoder). To prevent dehydration, 2ml of saline was administered subcutaneously every two hours. At the conclusion of each experiment, the animals were euthanized by isoflurane overdose, with exposure maintained for at least 1 min after cessation of respiration, in accordance with AVMA guidelines. The experimental sessions lasted 6–11 h, from anesthetic induction to euthanasia, with electrode implantation and recordings typically completed within 4–6 h.

### 2.3 High-Density epimysial EMG recordings

The fascia of the hindlimb was excised to reveal the gastrocnemius muscles, allowing placement of a 32-channel flexible high-density electrode array (“Rat EEG Triangular H32,” NeuroNexus Inc.) on the surface of the gastrocnemius lateralis. The array was positioned distally to proximally and aligned with the muscle innervation (Figure 1a). Monopolar EMG signals were acquired using a Nano2 headstage (Ripple Neuro, Inc) and sampled at 30 kS/s. All raw data were recorded and stored using a digital acquisition system (Scout, Ripple Neuro, Inc.) for subsequent offline analyses. Trigger signals were transmitted from the stimulator to the digital acquisition system concurrent with stimulation delivery, ensuring precise synchronization between stimulation events and HD-eEMG recordings, thereby supporting comprehensive post-processing and analysis.

### 2.4 Strength-Duration Curves and Stimulation Thresholds

The stimulation thresholds for each implanted electrode were determined from the strength-duration (SD) curves. Stimulation consisted of biphasic, square, cathodic-first pulses delivered at 1 Hz, utilizing either monopolar or bipolar configurations (Figure 1b). For each pulse width (PW) tested, ranging from 20µs to 300µs, the stimulation current (pulse amplitude, PA) was incrementally augmented until a visible muscle twitch was observed. The minimum current required to elicit a muscle twitch was defined as the activation threshold. This procedure was repeated across all electrodes for each stimulation strategy under evaluation. Electrodes that did not produce muscle twitches were excluded from further analysis.

### 2.5 Stimulation protocol - Amplitude Sweep

Rheobase was determined from the corresponding SD curves for each individual electrode or electrode combination. The stimulation pulse width was selected on the basis of the chronaxie value, as shown in Figure 1b. For each electrode or electrode combination, the baseline stimulation level (threshold) was defined as the pulse amplitude associated with the selected pulse width.

To assess the recruitment of muscle fibers via monopolar or bipolar stimulation, recruitment profiles were generated by incrementally adjusting the pulse amplitude in steps of 2µA or 4µA. This evaluation commenced at two increments below the activation threshold and continued up to 1.5 times the threshold value (xTH), as illustrated in Figure 1b. Stimulation consisted of biphasic, charge-balanced, square pulses in a monopolar or bipolar configuration (schematic shown in Figure 1b). For monopolar stimulation, cathodic first pulses were applied through a single LIFE, while for bipolar stimulation, two LIFEs were used, with one delivering cathodic-first pulses and the other delivering anodic-first pulses simultaneously. Stimulation was applied at 10 or 15 Hz in 5 s trains, with each train followed by a rest period of 20-30 seconds to minimize muscle fatigue. High-density EMG signals were continuously recorded throughout the stimulation protocol.

### 2.6 Data Processing

Monopolar EMG recordings were spatially filtered to decrease the detection volume and isolate the electrical activity of the underlying muscle, as described by Gallina et al. (28) (Figure 1c). A single-differential spatial filter was applied along the direction of the muscle fibers (columns) by subtracting each channel from its immediately adjacent channel in the column direction (i.e., Channel _(r,c)_ = Channel _(r,c)_ – Channel_(r+1,c)_, where r denotes the row and c denotes the column). Subsequently, temporal filtering was performed using a fourth-order Butterworth bandpass filter with cut-off frequencies of 10-1000 Hz.

For each stimulation intensity, stimulus-triggered averages (STA) of eEMG from each HD-eEMG channel were calculated to construct M-wave responses. To characterize these responses, the STA root mean square (RMS) amplitude of each M-wave was computed across a 16-ms window, beginning with the onset of the evoked response.

To assess the spatial distribution of EMG amplitudes across the electrode grid at each stimulation level, the RMS values for all EMG channels were normalized to the highest RMS value for the respective electrode configuration and stimulation intensity. These normalized RMS values were subsequently used to construct spatial heatmaps of stimulation-evoked muscle activity for visualization (Figure 1c). Next, the channels were sorted in the descending order of their normalized RMS values, which were then plotted against the sorted channel order. This created an amplitude rank profile that illustrated the distribution of the RMS values among the channels. The area under the curve (AUC) for each profile curve, corresponding to each stimulation level for each rat and stimulation configuration, was calculated using trapezoidal numerical integration (MATLAB, *trapz*). Lower AUC values indicate that the EMG amplitude was concentrated in fewer channels, reflecting greater selectivity, whereas higher AUC values denote a more evenly distributed EMG amplitude across channels, indicating reduced selectivity.

### 2.7 Statistical Analyses

All statistical analyses were conducted using R (version 4.5.0, R Core Team, 2024), with statistical significance set at α = 0.05. Linear mixed-effects regression models were employed to address our research questions, with the fixed and random effects detailed below. These models were implemented using the *lme4* package (31) utilizing Restricted Maximum Likelihood Estimation. Where applicable, post hoc comparisons were performed using estimated marginal means calculated with the *emmeans* package (32), with adjustments for multiple comparisons (Tukey or Dunnett). Model assumptions, including linearity and normality of residuals, were evaluated to ensure the validity of model fitting. The model equations are presented in R syntax rather than in standard linear mixed model notation. Prior to fitting the statistical model, the stimulation level was mean-centered to facilitate the interpretation of the regression coefficients. This approach ensured that the model intercept represented the eEMG amplitude response at the mean stimulation level, rather than at a stimulation level of zero.

#### 2.7.1 Effect of stimulation level on EMG amplitude

To assess the effect of monopolar Stimulation Amplitude level on EMG amplitude across channels, a linear mixed-effects model was constructed with fixed effects of *Stimulation Amplitude*_*c*_ *(centered), EMG Channel*, and *Stimulation Amplitude*_*c*_*-by-EMG Channel* interaction. The dependent variable was the RMS amplitude of the M-wave response (including data from all EMG channels). To fully account for dependencies within the data, we included random intercepts for *Rat, Channel* (as a repeated sampling of muscle locations within each rat) within *Rat*, and *LIFEs* within *Rat*, and random slopes for *Stimulation Amplitude*_*c*_ within *LIFEs* within *Rat*.

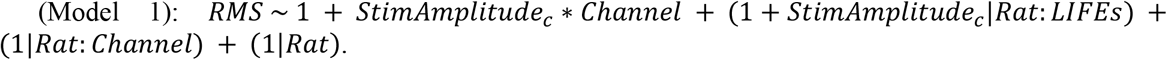

#### 2.7.2 Effect of stimulation from different LIFEs on EMG amplitude

The model from the previous section (the effect of *Stimulation Amplitude*_*c*_*-by-EMG Channel* on RMS amplitude) was used to assess the effect of different LIFEs delivering monopolar stimulation on EMG amplitude across channels and at different stimulation amplitudes. The experimental design precluded the inclusion of *LIFEs* as a fixed effect, given that LIFE placement was not anatomically standardized across animals; LIFE #1 in one rat was not implanted at a location equivalent to LIFE #1 in another rat. Therefore, the variance for a unique electrode (the random intercept of *LIFEs* within *Rat*) and the variance of the effect of Stimulation Amplitude for a unique electrode (the random slope for *Stimulation Amplitude*_*c*_ by *LIFEs* within *Rat*) were examined in relation to the global average RMS value and the variance due to *Rat* and *Channel* within Rats.

1. Effect of stimulation from different LIFEs on EMG spatial distribution

To assess whether spatial selectivity (quantified as the area under the curve (*AUC*) of ranked EMG amplitude profiles varied with stimulation level, a linear mixed effects model was constructed with a fixed effect of *Stimulation Amplitude*_*c*_ and a random intercept for *Rat*, and a random slope for *Stimulation Amplitude*_*c*_ by *LIFE* within *Rat*.

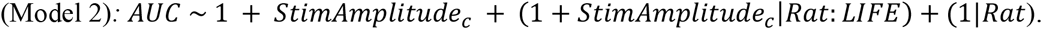

The shape of the ranked amplitude profiles at each stimulation level was averaged across all electrodes, and the resulting curve was characterized using polynomial regression to investigate how the spatial distribution of EMG changed with stimulation intensity. For each stimulation level, the mean-ranked amplitude profile was fitted using least squares to a cubic polynomial in the channel rank in the following form:

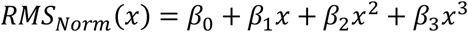

where *β*_0_ is the intercept term, *β*_1_-*β*_3_ are the coefficients of the polynomial curve, and *x* is the ranked channel. The fitted coefficients were used to assess how the profile shape changed with increasing stimulation levels.

The standard deviation was calculated for the mean profile across rats per stimulation level to examine the variability of the ranked amplitude profiles across rats. This resulted in an SD profile, a calculated mean SD, and the area under the SD Curve (*trapz*, MATLAB).

#### 2.7.3 Effect of stimulation type on EMG amplitude

To evaluate the effect of stimulation type (monopolar or bipolar) on EMG amplitude across channels and at different stimulation levels, a linear mixed effects model was constructed with fixed effects of *Stimulation Type, Stimulation Amplitude*_*c*_, *EMG Channel*, and a three-way interaction of *Stimulation Type-by-EMG Channel-by-Stimulation Amplitude*_*c*_.

The *Stimulation Type* encompassed Monopolar and Bipolar modalities. In the bipolar stimulation configuration, the electrodes were labeled solely as MS1 and MS2 to distinguish their polarity assignment; both electrodes were activated simultaneously with MS2 having its polarity reversed with an anodic first pulse. This labeling carried no distinction in the monopolar stimulation configuration, where the cathodic first pulses were delivered to both electrodes. The dependent variable was the RMS amplitude of the M-wave response. To fully account for dependencies within the data, we included random intercepts for *Channel* within *Rat*, and a random slope for *Stimulation Amplitude*_*c*_ by *Stimulation Type* within *Rat*.

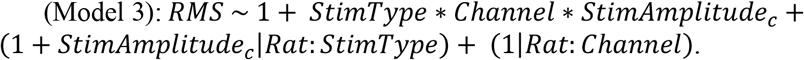

#### 2.7.4 Effect of stimulation type on eEMG spatial distribution

To assess whether spatial selectivity (quantified as the area under the curve (*AUC*) of ranked eEMG amplitude profiles) varied among the three stimulation types, a linear mixed effects model was constructed with fixed main effects of *Stimulation Type* and *Stimulation Amplitude*_*c*_, *and the Stimulation Type-by-Stimulation Amplitude* interaction. We included a random intercept for *Rat* and a random slope for *Stimulation Amplitude*_*c*_ by *Stimulation Type* within *Rat*.

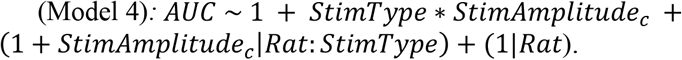

#### 2.7.5 Assessment of spatial localization of highest HD-eEMG channel activation across stimulation levels

For each stimulation level, the channel showing the maximum value for each electrode was identified, and the number of occurrences per channel was counted to build histograms and perform the analyses. Analyses were performed separately for the two types of stimulation (monopolar and bipolar), at each level.

## 3. Results

We collected M-wave responses on stimulation using 49 LIFEs across 17 different rats using a monopolar stimulation configuration and from 27 pairs of electrodes using a bipolar stimulation configuration in a subset of nine rats.

### 3.1 Monopolar stimulation induces graded muscle activation

Figure 2 illustrates the effects of increasing the stimulation level using four different LIFEs with M-wave responses from the individual eEMG channels. Monopolar stimulation resulted in graded muscle activation, which increased with the increasing stimulation amplitude. There was a significant main effect of *Stimulation Amplitude* (F(1,48) = 39, *p* < 0.0001) and a significant *Stimulation Amplitude-by-Channel* interaction (F(25,17457) = 21, *p* < 0.0001), indicating that the effect of increasing the stimulation amplitude on RMS differed among the HD-eEMG channels.

**Figure 2.**
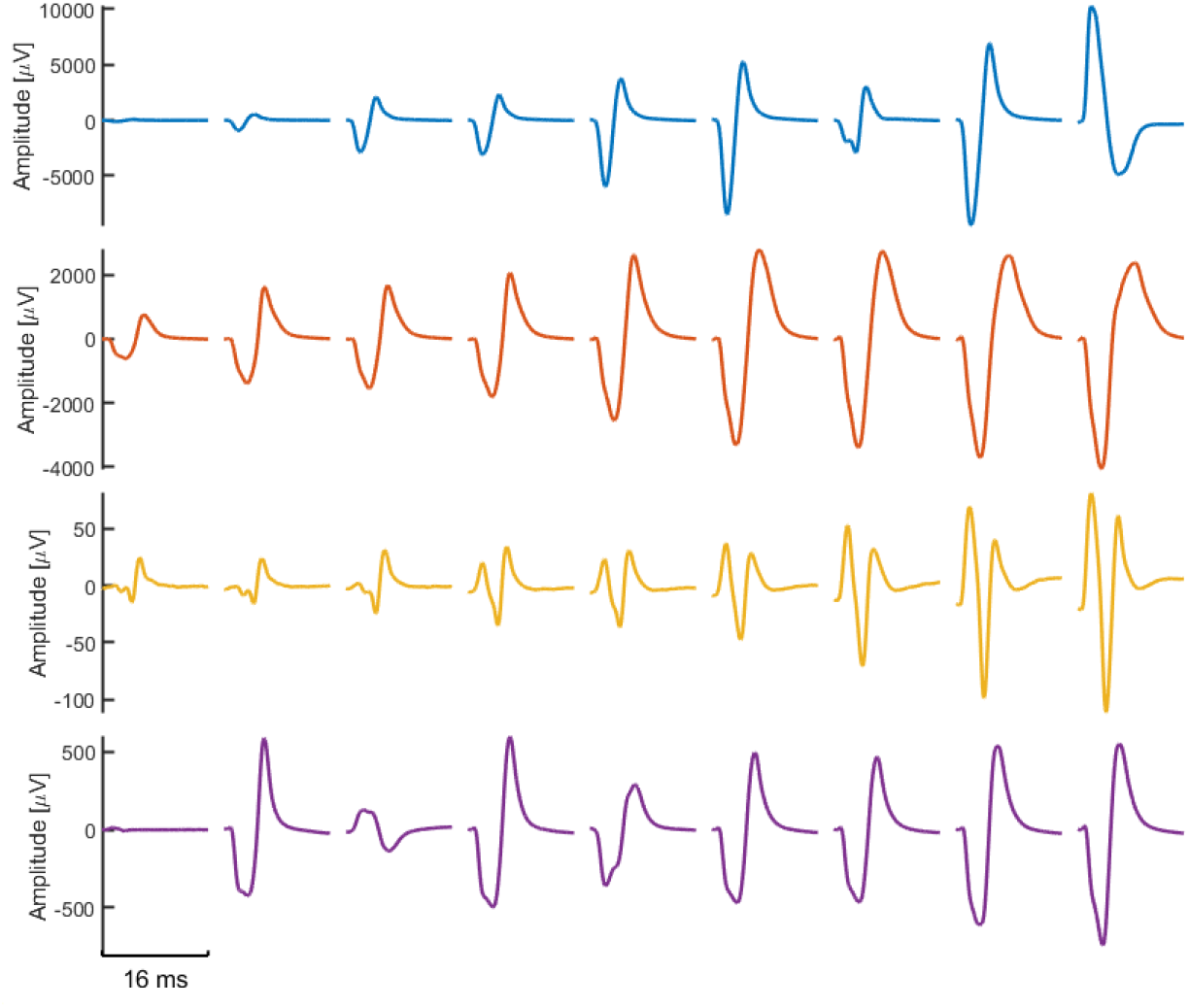
Comparison of M-wave responses to monopolar stimulation using four LIFEs across the amplitude sweep. The effect of increasing stimulation level on M-wave responses recorded from one HD-eEMG channel for four different LIFEs are shown. Each trace for a given row is the stimulus-triggered average response amplitude (y-axis) over time (x-axis – 16ms each segment) for nine stimulation levels (increasing in amplitude from left to right) and illustrates graded activation. Stimulation amplitudes are normalized to the activation threshold for each individual LIFE. The starting stimulation amplitude was set below this threshold to capture the progressive increase in neural recruitment.

The nature of the graded activation differed slightly depending on the LIFE used. Simulations with LIFE #1 elicited subtle changes in M-wave shape at the initial stimulation level, with more pronounced responses after the fourth stimulation level. The increase in M-wave amplitude was generally monotonic, with one abrupt reduction at the seventh stimulation level. The highest stimulation level resulted in the reversal of the M-wave polarity. Stimulation with LIFE #2 and LIFE #3 showed more consistent graded increases in response magnitude across stimulation levels. Finally, stimulation with LIFE #4 demonstrated a less consistent increase at lower stimulation levels, with a subtle but consistent increase in response magnitude at the five highest stimulation levels.

### 3.2 Stimulation using multiple LIFEs implanted in the same fascicle produces variable response magnitudes and different spatial patterns of muscle activity

#### 3.2.1 Effects on muscle response magnitude

We examined the standard deviation associated with random intercepts of *Rat, LIFE* (nested within Rat), and *Channel* (nested within *Rat*) and the random slope of *Stimulation Amplitude* by *LIFE* (nested within *Rat*) from our *Stimulation Amplitude*-by-*Channel* model, to assess their relative influences on the variability of RMS response magnitude.

The estimated global mean RMS value of the M-wave responses across rats, LIFEs, eEMG channels, and stimulation amplitudes was 428.7 µV. The standard deviations due to each random effect are listed in Table 1. The random effect of LIFE contributed the greatest variability to the RMS, with a standard deviation of 405.4 µV. This indicates that while the mean RMS value was 428.7µV, for any given LIFE, the RMS could vary around 428.7 with a standard deviation of +/-405.4 µV. The standard deviations of the RMS value due to *Rat* and *Channel* were lower, at 290.4 and 251.6 µV, respectively. The substantial variability in the RMS values of the responses obtained using different LIFEs demonstrates their flexibility and potential for further development of targeted selectivity.

**Table 1.**
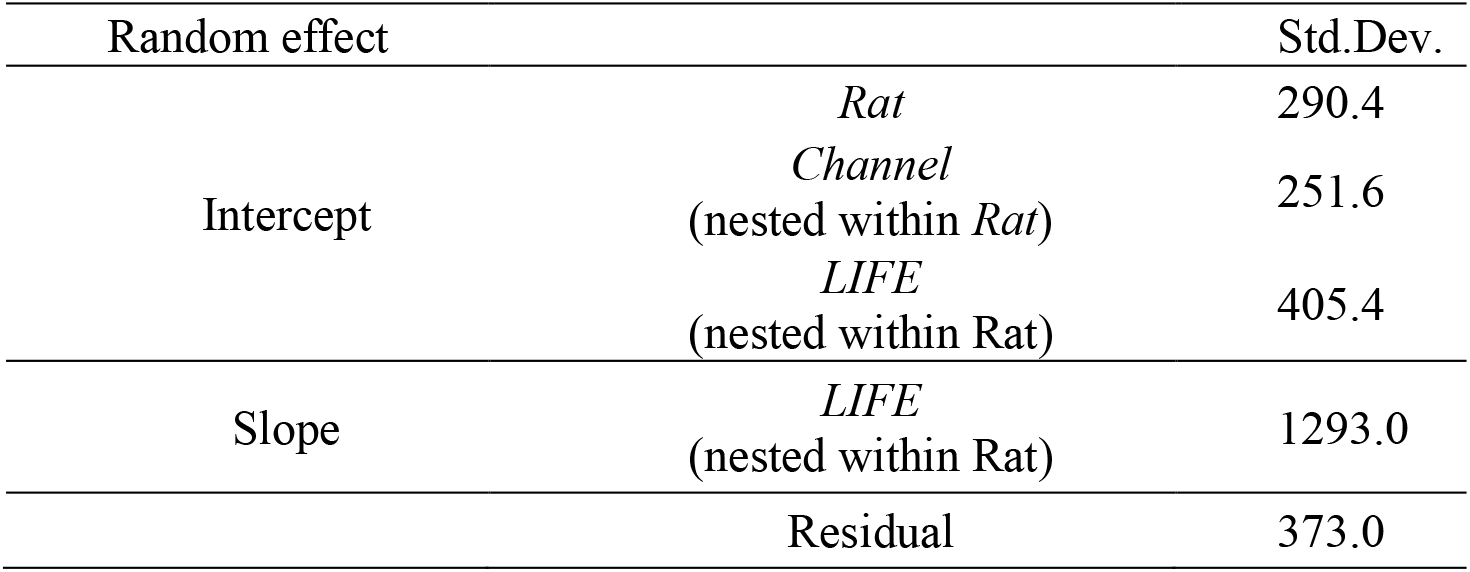
Standard deviation in RMS value attributed to each random effect from the Stimulation Amplitude-by-Channel model.

The effect of the *Stimulation Amplitude*_*c*_ also varied substantially among LIFEs. The model estimates for the effect of *Stimulation Amplitude*_*c*_ (on average among LIFEs) was 1298.9 µV, indicating the amount that RMS increased per 1 threshold level increase in stimulation level. This slope varied among LIFEs with a standard deviation of 1293 µV, demonstrating that the rate of RMS increase with increasing stimulation intensity is highly dependent on the specific LIFE used for stimulation; the variability is almost equal in magnitude to the average effect.

#### 3.2.2 Effects on spatial muscle activation patterns

We evaluated the variation in spatial muscle activation during stimulation across different LIFEs. Upon analyzing a single HD-eEMG array channel (Figure 3a, circled in green), the corresponding M-wave (Figure 3b) displayed distinctly different waveform shapes, depending on which LIFE was utilized for stimulation. This observation implies that stimulation using each LIFE recruited a different combination of nerve fibers. Further assessment of the entire HD-eEMG grid revealed that RMS amplitude heatmaps demonstrated pronounced differences in the spatial distribution of muscle activation among the LIFEs, including the channel that exhibited the highest amplitude. Representative data from one rat are shown in Figure 3. Among the LIFEs used, the activation patterns produced by LIFE #2 and LIFE #4 were the most similar, exhibiting a peak activation along the left side of the grid. Conversely, the use of LIFE #3 produced a markedly distinct pattern with peak activation observed on the right side of the grid. Stimulation with LIFE #1 resulted in a broader activation pattern across the entire grid.

**Figure 3.**
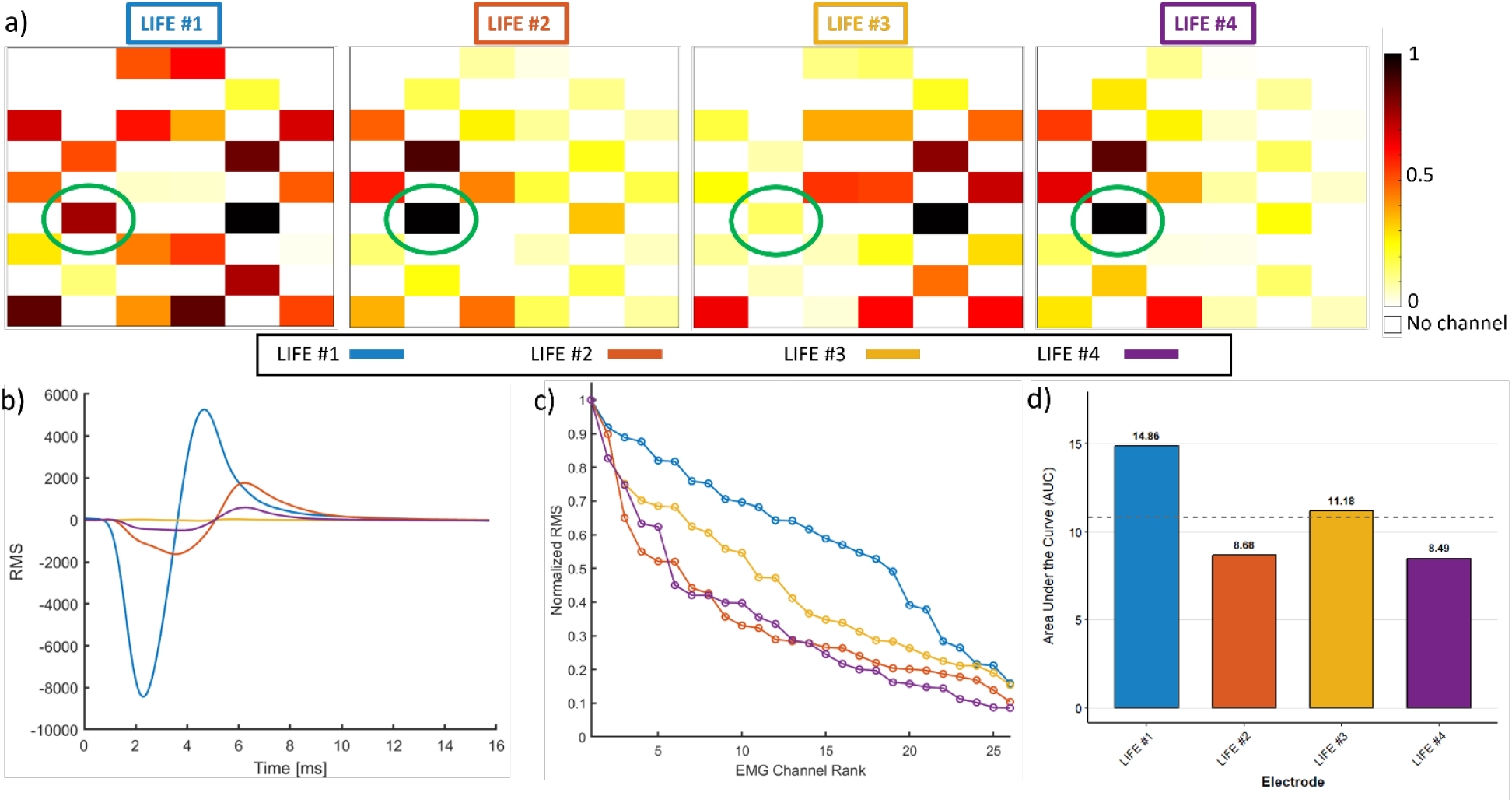
Comparison of spatial muscle activation resulting from monopolar stimulation configurations. (a) Heatmaps showing eEMG amplitude across channels for stimulation using four different LIFEs (LIFE #1 - #4) in one rat at one stimulation level (1.2xTH), with color representing Root Mean Square (RMS) amplitude, normalized to the highest value across all eEMG Channels for each grid. Green circles highlight the spatial location of the highest muscle activation obtained across the grid when using LIFE #2 or LIFE #4, demonstrating that each LIFE evokes distinct muscle activation patterns. (b) Stimulus-Triggered Averages (STA) M-wave responses for each LIFE at the same stimulation level (xTH) showing different magnitude and waveform shapes. (c) Ranked normalized RMS profile curves across EMG Channels, demonstrating that stimulation using LIFE #1 elicited the broadest activation with low selectivity, while use of LIFE #2 and LIFE #4 exhibited higher selectivity with lower diffusivity of activation. (d) Bar plots showing the calculated area under the curve (AUC) from the ranked RMS profile curves.

The ranked eEMG amplitude profiles corresponding to the same data are shown in Figure 3c, and their respective AUC values are shown in Figure 3d. In alignment with the visual assessment of the RMS value heatmaps, the use of LIFE #1 demonstrated the least selective recruitment, as evidenced by the relatively flat profile curve and the highest AUC. In contrast, LIFE #2 and LIFE #4 exhibited steeper profile curves and lower AUC values, reflecting a more spatially focused activation concentrated within fewer eEMG channels. Stimulation using LIFE #3 showed an intermediate profile, occupying a position between these two extremes.

#### 3.2.3 Effects of stimulation amplitude and choice of LIFEs on spatial activation

Figure 4 shows the effect of the stimulation amplitude on the distribution of eEMG and how it varies among the LIFEs chosen for stimulation. Figure 4a shows data from the same LIFE as Figure 3, demonstrating that the AUC of the ranked eEMG amplitude profiles remained relatively stable across stimulation levels, with minor variations observed only for stimulation using LIFE #1 and LIFE #4.

**Figure 4.**
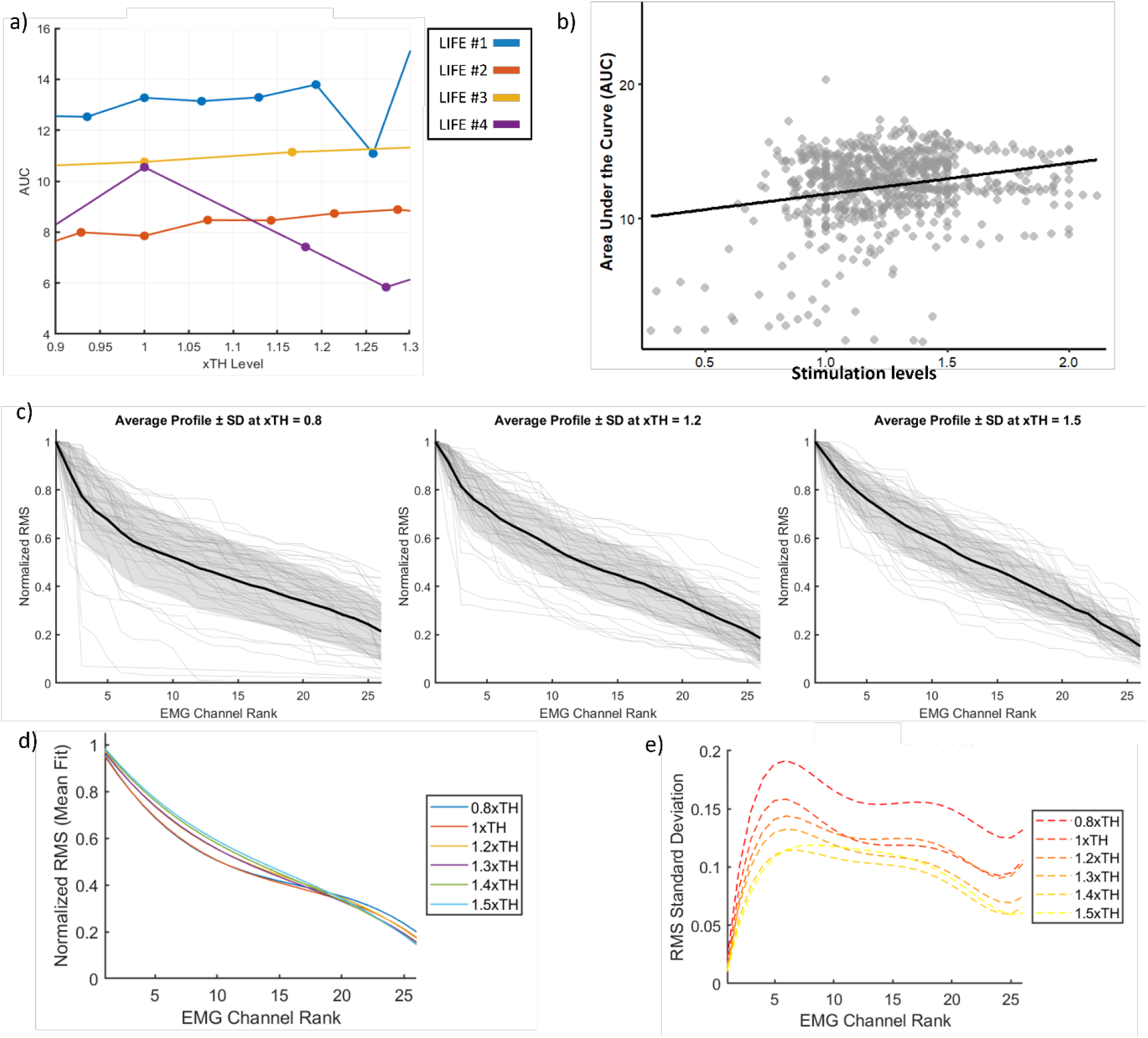
Quantification of intrafascicular activation on stimulation using different LIFEs across stimulation levels. (a) Example from a single rat showing the area under the curve (AUC) for stimulation using four different LIFEs across different stimulation levels (same animal as from Figure 2). (b) Relationship between AUC and normalized stimulation threshold. Each point represents data for an individual LIFE at that stimulation level. The solid black line shows the linear mixed model. (c) Ranked normalized RMS profile curves of all electrodes (grey) at three different stimulation levels: subthreshold (0.8xTH), slightly above threshold (1.2xTH) and highest stimulation level (1.5xTH). The average profile is shown in black with its standard deviation in shaded gray. (d) Polynomial curves fit to the average ranked RMS profiles for each simulation amplitude bin. (e) The standard deviation around the average ranked eEMG amplitude as a function of stimulation levels (xTH).

Analysis of the complete dataset using the linear mixed-effects model (Model 2) demonstrated a significant effect of stimulation amplitude on the AUC (*F*(1, 46) = 15.0, *p* < 0.001) (Figure 4b). This finding indicates that higher stimulation levels are associated with increased muscle activation among eEMG channels, resulting in higher AUC values and a concomitant decrease in selectivity.

Nevertheless, the magnitude of this effect (an estimated increase of 2.3 AUC for a 1x threshold increase in stimulation) was small, indicating that stimulation intensity had a limited effect on selectivity, as did the model’s pseudo-*R*^*2*^ (0.05), indicating that stimulation amplitude accounted for only a limited proportion of the variance in AUC.

The random intercept variance associated with electrodes nested within rats suggested modest but significant differences in the AUC across electrodes (variance = 4.27, SD = 2.06), highlighting the contribution of electrode position to the variability in recruitment and selectivity. This variance was higher than that attributed to *Rat* alone (SD = 0.88) and random effects (SD = 1.59). For more context, the mean AUC across all data was 12.39 a.u., hence the electrode-level variance accounted for almost 18% of the mean AUC. These findings indicate the importance of electrode placement in influencing the recruitment and selectivity.

Figure 4c shows the ranked eEMG amplitude profiles derived from the full dataset at three different stimulation amplitudes. Individual profiles for each LIFE used are depicted in gray, wheras the mean profile and its standard deviation are shown in black and shaded gray, respectively. The overall shape of the profiles gradually changed across the stimulation levels, becoming progressively flatter and more linear as the stimulation amplitude increased. The coefficients obtained from the polynomial fit of the mean profile for each stimulation level are listed in Table 2. As the stimulation amplitude increased, both the negative cubic and linear coefficients exhibited an upward trend, the positive quadratic coefficient exhibited a downward trend, and the intercept remained stable. This systematic pattern is consistent with broader activation of the muscle and reduced selectivity with increasing stimulation amplitude.

**Table 2.** Coefficient of polynomial fit for profile curves at each stimulation level (Monopolar Stimulation)

| xTH | $\beta_3 (x 10^{-5})$ | $\beta_2 (x 10^{-3})$ | $\beta_1 (x 10^{-2})$ | $\beta_0$ |
| --- | --- | --- | --- | --- |
| 0.8 | -9.9 | 4.9 | -9.3 | 1.04 |
| 1 | -9.2 | 4.5 | -8.9 | 1.04 |
| 1.2 | -7.5 | 3.7 | -7.9 | 1.05 |
| 1.3 | -6.6 | 3.3 | -7.4 | 1.04 |
| 1.4 | -5.8 | 2.9 | -6.9 | 1.04 |
| 1.5 | -6.0 | 2.9 | -6.9 | 1.05 |

In addition to the changes in the average profile shape as a function of stimulation intensity (Figure 4d), notable differences in the variability of the individual profiles were also apparent (Figure 4e). At sub-threshold stimulation levels (0.8 xTH), individual profile curves displayed considerable variability among the LIFEs used. This variability qualitatively decreased at 1.2 xTH and reached its minimum at 1.5 xTH. This pattern indicated a more consistent pattern of muscle activation across the electrodes and animals as the stimulation amplitude increased.

We also cataloged the channel with the maximum RMS amplitude for each LIFE used, and the stimulation amplitude condition for all conditions. Figure 5a shows histograms of the frequency of occurrence for each channel. For all stimulation amplitudes, the frequency distribution of channels with the maximum RMS value was not normal, with multiple peaks (Figure 5a), indicating a non-random spatial organization of muscle activation. Instead, the distributions were concentrated in a limited set of channels that were predominantly located near the center of the grid. They were also clustered into distinct regions, showing off-center peaks (Figure 5b). Although increasing the stimulation amplitude produced modest shifts in the relative frequency of occurrence, the locations of the peaks generally remained stable (channels 8, 6, and 21).

**Figure 5.**
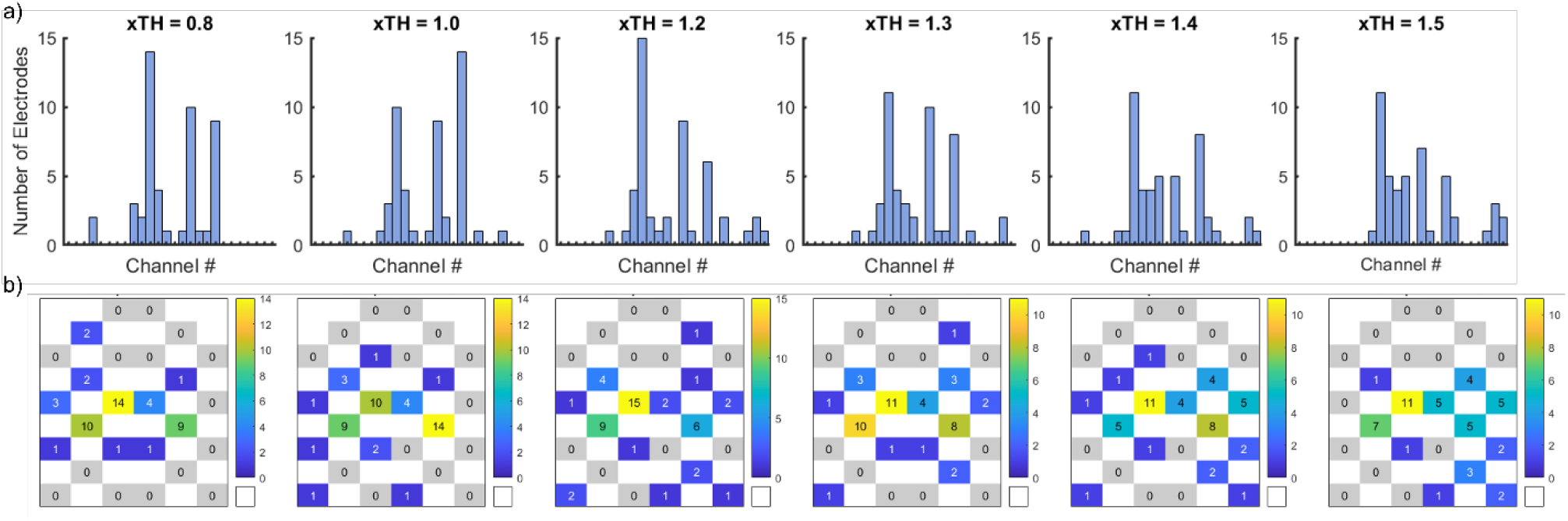
Assessment of distribution of highest value EMG channel across all LIFEs. (a) Histograms showing the EMG channel with the highest value of the grid across different stimulation levels (xTH). (b) Heatmaps from the histograms in (a) showing the spatial distribution of the most activated EMG channel position on the grid across all stimulation levels. A concentrated heatmap indicates a consistent highest EMG channel position, whereas a diffuse distribution reflects variability in the location of peak activation on the muscle across stimulation levels.

Figure 6 presents a comparison of the graded M-wave responses across monopolar and bipolar stimulation configurations. The top and middle rows illustrate the responses obtained from monopolar stimulation using two distinct LIFEs, designated Monopolar - LIFE #3 and Monopolar – LIFE #2 respectively. The bottom row depicts the responses from bipolar stimulation in which the same two LIFEs were utilized in a paired configuration. Across both monopolar and bipolar stimulation configurations, the magnitude of the evoked responses increased as the stimulation amplitude increased. This demonstrates that both simulation configurations exhibit a graded recruitment of muscle fibers, with higher amplitudes producing greater M-wave responses.

**Figure 6.**
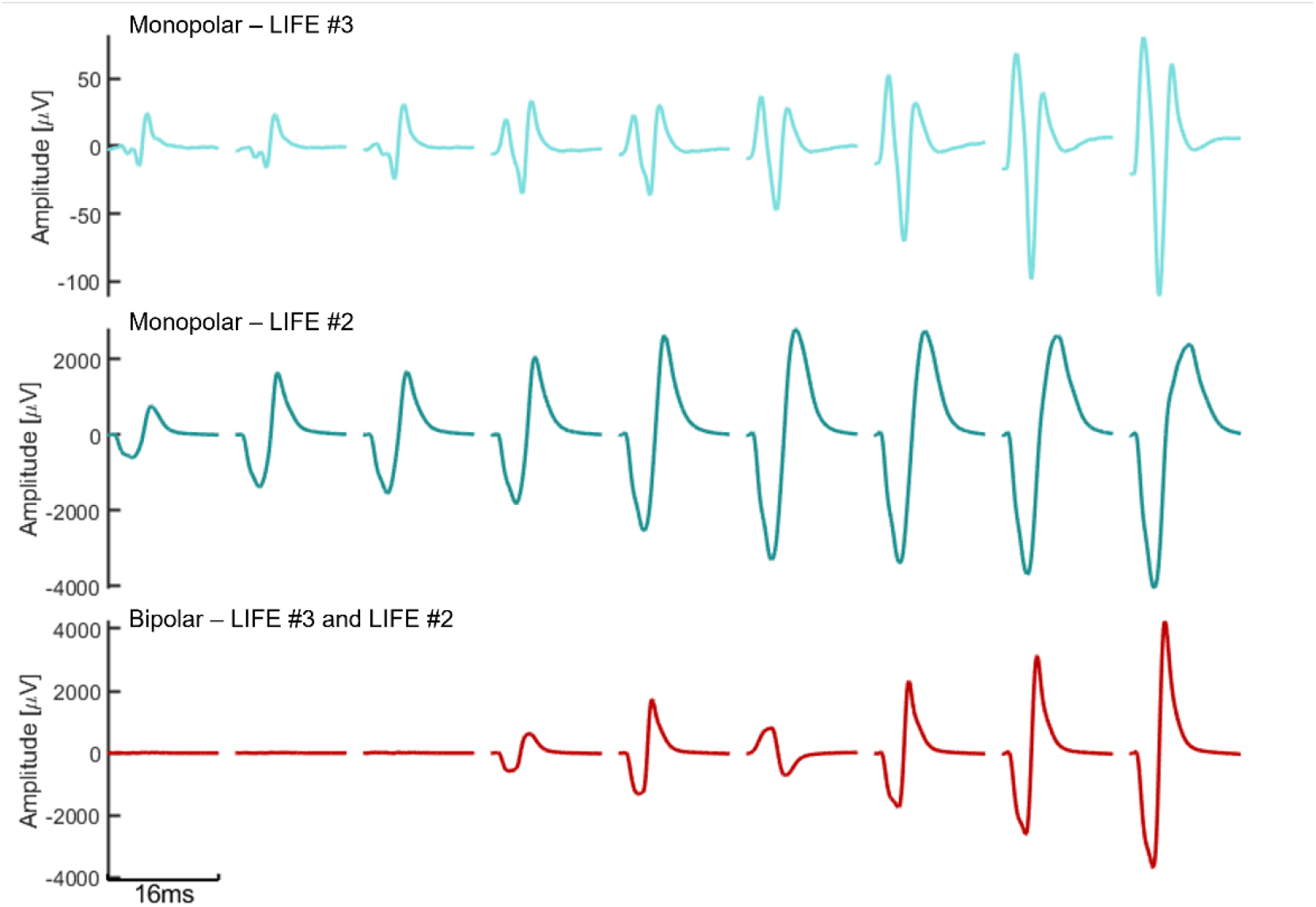
Comparison of graded M-wave responses across monopolar and bipolar stimulation configurations. The magnitude of evoked M-wave responses increased with increasing stimulation amplitude. Top and middle rows: example responses from one rat on monopolar stimulation using LIFE #3 (Light blue; top) and LIFE#2 (Dark Blue; middle). Bottom row: example responses on bipolar stimulation using both LIFE #3 and LIFE #2 (red). Each trace for a given row is the stimulus-triggered average response amplitude (y-axis) over time (x-axis of 16ms width) for nine stimulation levels (increasing in amplitude from left to right) and illustrates graded activation.

#### 3.2.4 Bipolar and monopolar stimulation produce different muscle activation patterns

As illustrated in Figure **7**, bipolar stimulation utilizing two LIFEs resulted in HD-eEMG array heatmaps (a), M-wave responses (b), and ranked eEMG amplitude profiles (c, d) that exhibited differences compared to the outcomes of separate monopolar stimulations with the same electrodes. In this representative example, the spatial activation pattern generated using LIFE #2 in a monopolar configuration closely resembled that observed with bipolar stimulation, whereas monopolar stimulation using LIFE #3 produced a significantly different, nearly inverted muscle activation pattern. Analysis of the ranked eEMG amplitude profiles revealed that monopolar stimulation using LIFE #2 yielded the steepest curve and the lowest AUC, indicating the highest selectivity among the three configurations. Bipolar stimulation produced a similar profile, although with slightly reduced selectivity, whereas stimulation using LIFE #3 resulted in broadest spatial activation.

**Figure 7.**
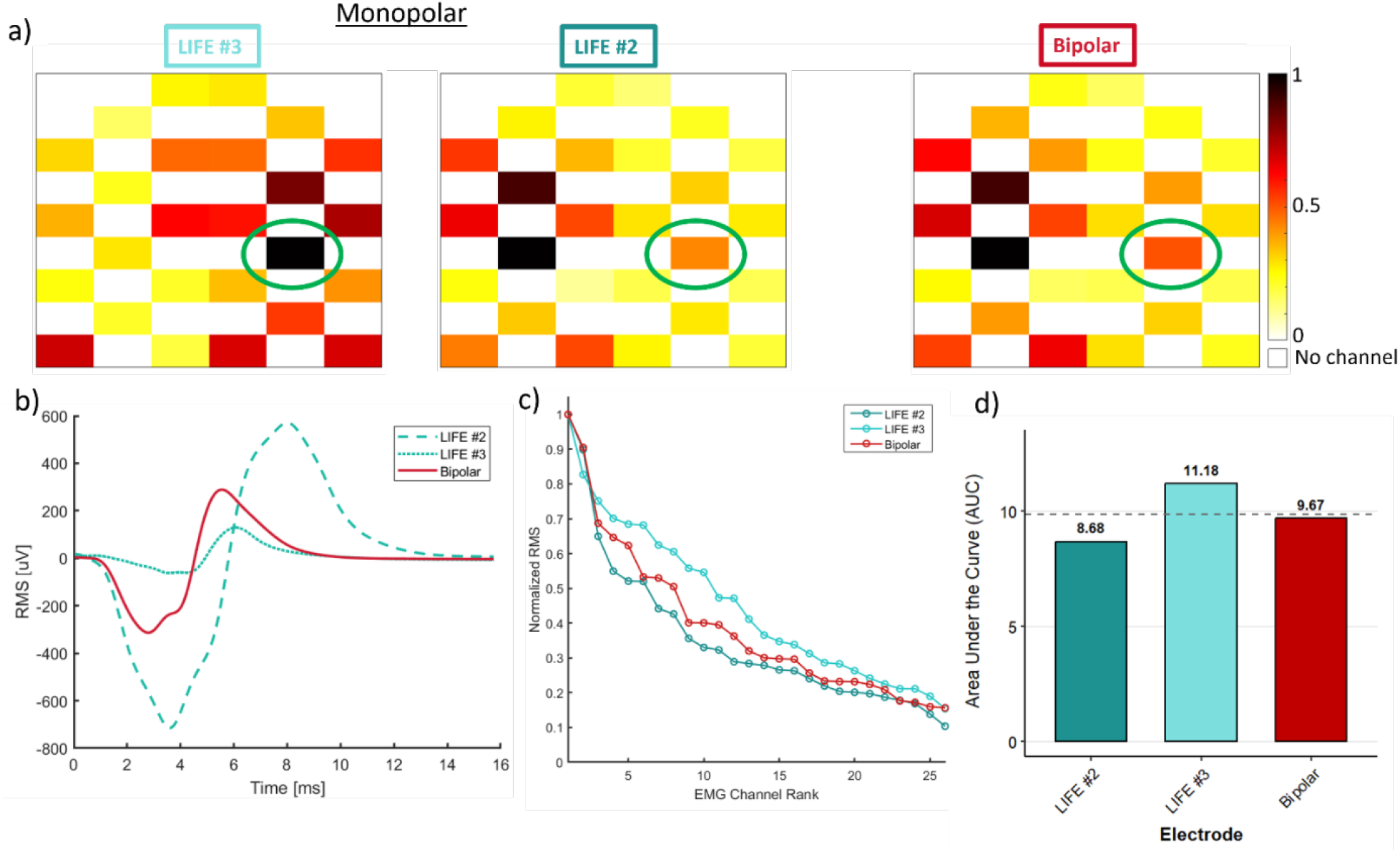
Comparison of spatial HD-eEMG elicited on monopolar vs. bipolar stimulation configurations. Data from a representative rat. (a) Heatmaps show HD-eEMG for all channels resulting from monopolar stimulation with two separate LIFEs (Monopolar - LIFE #3 (left), Monopolar - LIFE #2 (right)) and bipolar stimulation, with color representing normalized Root Mean Square (RMS) amplitude. Green circles highlight the spatial location of the highest muscle activation for stimulation using LIFE #3. (b) Representative Stimulus Triggered Average M-wave response from the HD-eEMG channel circled in green in (a), for the two monopolar and the bipolar stimulation configurations. Each stimulation was performed at the same physiological level (xTH) but resulted in different response magnitudes and waveform shapes. (c) Ranked RMS profile curves for the two monopolar and the bipolar stimulation configurations. (d) Area Under the Curve (AUC) from the ranked RMS profile curves, with higher values indicating broader spatial activation and lower selectivity.

Although Figure **7** is from a single rat, there was a significant *StimulationType* x *Channel* interaction when evaluating the entire dataset statistically (F(50, 9717) = 8.6), *p* < 0.0001), which is consistent with the patterns visualized in the example.

#### 3.2.5 Effect of increasing stimulation amplitude on spatial EMG with bipolar vs. monopolar stimulation

There was a significant three-way *StimulationAmplitude*_*c*_ x *StimulationType* x *Channel* interaction (F(50, 9694) = 14.3), *p* < 0.0001), indicating that the effect of the stimulation amplitude on the eEMG location (across channels) depended on the stimulation type.

The size of the spread of eEMG (rather than the location), as assessed by the AUC of the ranked eEMG amplitude profile, was largely stable with increasing stimulation amplitude with bipolar stimulation, similar to that with monopolar stimulation (Figure 8a, b). In addition, the average ranked eEMG profiles for the monopolar and bipolar configurations showed no major qualitative differences in overall shape, steepness, or variability for any of the stimulation levels (Figure 8c, d, e). The coefficients of the polynomial fit for bipolar stimulation (not reported for brevity) were virtually identical to those for monopolar stimulation (Table **2**). Accordingly, there was no significant main effect of *StimulationType* on the AUC (*p* = 0.89; estimated means: monopolar MS1: 12.9, monopolar MS2: 13.2, and bipolar: 13.0), with no significant *StimulationType* x *StimulationAmplitude* interaction (*p* = 0.91).

**Figure 8.**
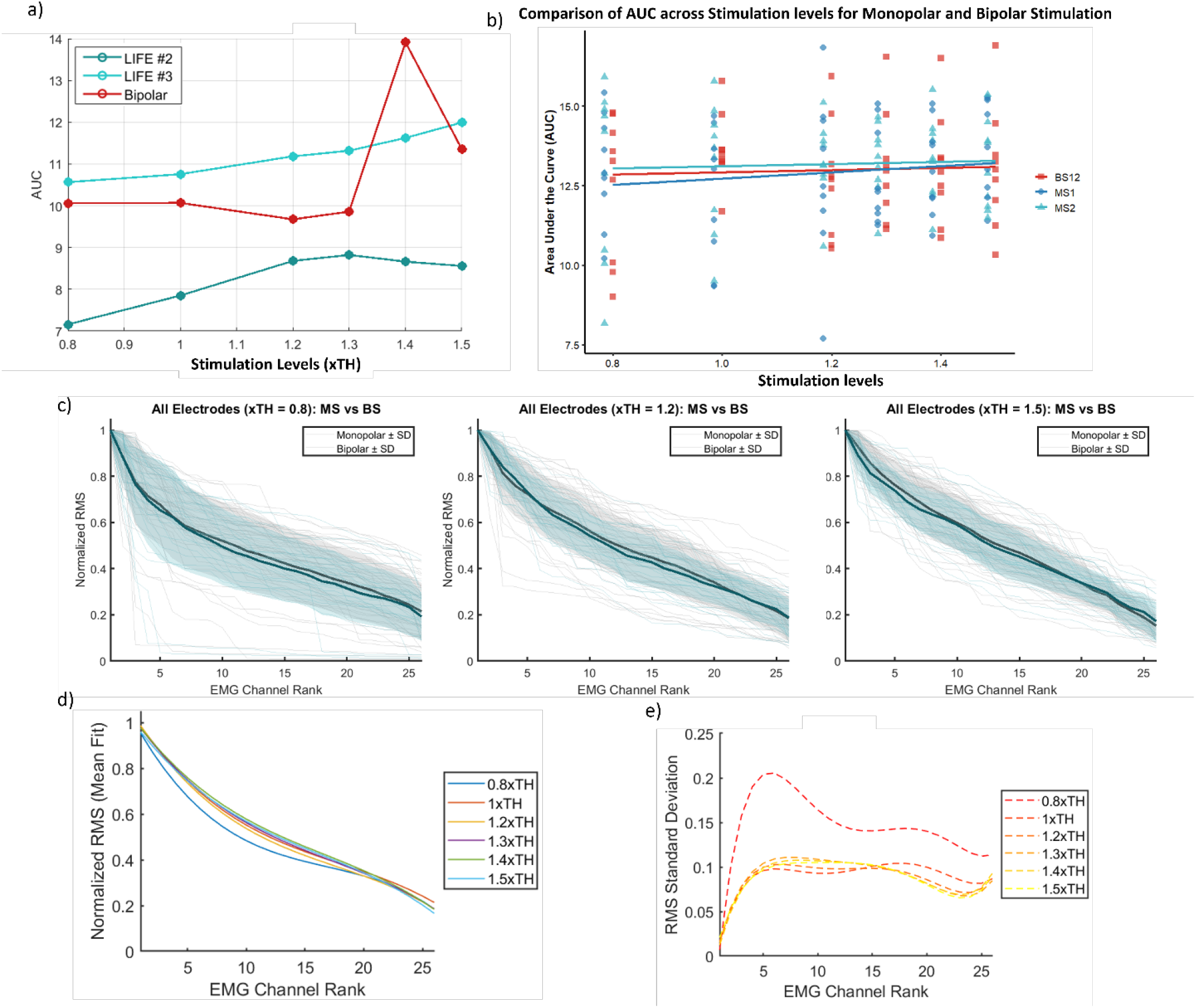
Assessment of intrafascicular selectivity for monopolar and bipolar stimulation configurations across stimulation levels. (a) Example showing the Area Under the Curve (AUC) for stimulation using two LIFEs in monopolar configuration and when used together in a bipolar configuration across different stimulation levels (same LIFEs as in Figure 6). (b) AUC of ranked eEMG amplitude profiles across all LIFEs and animals for each stimulation level and stimulation type (Monopolar configuration: Green, Bipolar configuration: Red). (c) Ranked eEMG amplitude profiles of all electrodes in monopolar configuration and pairs of electrodes in bipolar stimulation at three different stimulation levels: subthreshold (0.8xTH), slightly above threshold (1.2xTH) and highest stimulation level (1.5xTH). (d) Curves fitted to the mean normalized Root Mean Square (RMS) for bipolar stimulation across all the electrodes and stimulation levels. (e) Standard deviation of normalized RMS across ranked EMG channels for bipolar stimulation as a function of the stimulation level (xTH).

These findings indicate that while the different stimulation types altered the location of the eEMG and the effect of the stimulation amplitude on each channel, the size of the spread of the eEMG was not affected.

**Figure 9** compares the frequency distributions of the channel exhibiting the highest RMS value with bipolar versus monopolar stimulation. Both stimulation types had similar distributions, with multiple peaks (Figure 9a). However, compared with monopolar stimulation, the bipolar configuration showed a more concentrated distribution across fewer channels, as shown in Figure 9b, indicating that the channel with peak RMS values was more consistent than that with monopolar stimulation.

## 4. Discussion

In this study, we examined how to obtain a more selective peripheral nerve stimulation by activating the intended nerve fiber subpopulations while reducing off-target activation when using LIFEs. Two potential strategies were examined: electrode placement within the same fascicle and intrafascicular field steering. High-density epimysial EMG was used to quantify and map muscle activation patterns, providing a method to experimentally measure intrafascicular selectivity. Stimulation using LIFEs placed at different cross-sectional/longitudinal locations in the same fascicle can activate different muscle regions, indicating intrafascicular selectivity. Bipolar stimulation configurations can recruit fibers differently than monopolar configurations, offering a “lever” to shape the activated subpopulations. Both monopolar and bipolar stimulation evoked graded eEMG responses. Improving on-target selectivity is a path toward next-generation bioelectronic therapies with fewer side effects and better outcomes, potentially enabling more precise organ-specific neuromodulation.

**Figure 9.**
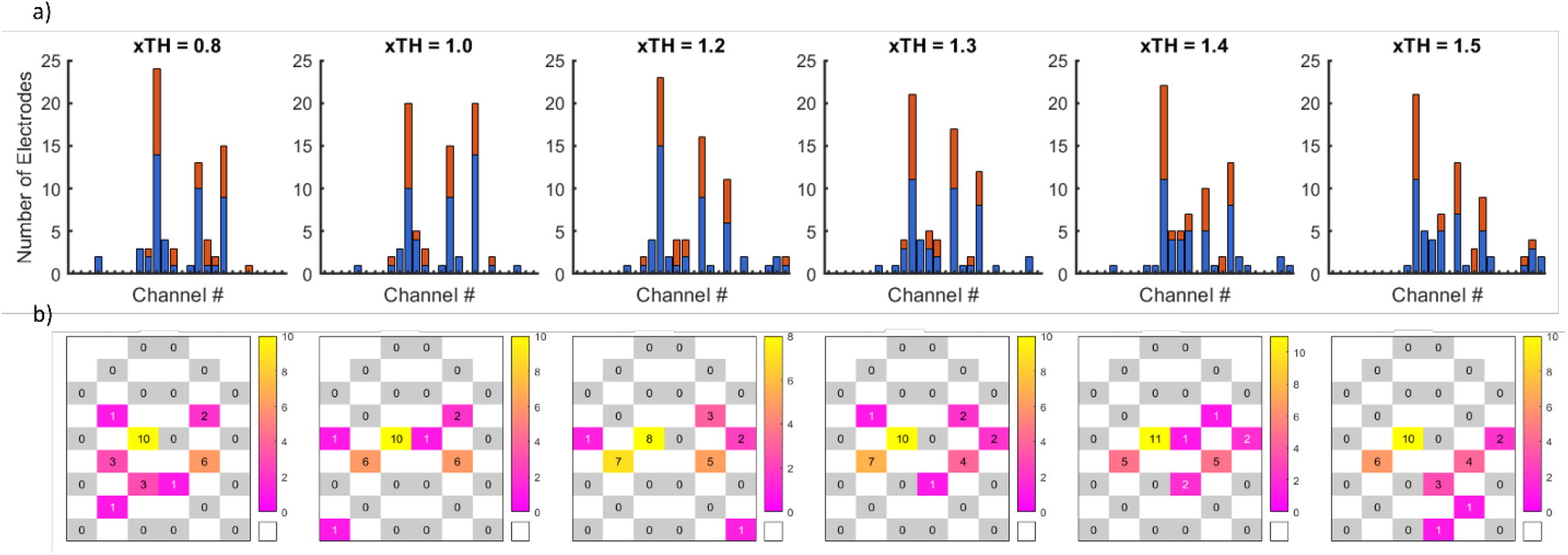
Assessment of distribution of highest value EMG channel across all LIFEs for both stimulation configuration. (a) Histograms showing the EMG channel number with the highest value of the grid across different stimulation levels (xTH) for bipolar stimulation (red) and monopolar stimulation (blue). (b) Heatmaps from the histograms showing the position on the grid of the highest value distribution across all stimulation levels (only bipolar stimulation is shown). A concentrated heatmap indicates a consistent highest EMG channel position, whereas a diffuse distribution reflects variability in the location of peak activation of the muscle across stimulation levels.

### 4.1 HD-eEMG enables detailed assessment of intrafascicular selectivity

High-density epymisial EMG is a practical, minimally invasive method for assessing intrafascicular selectivity during peripheral nerve stimulation. Unlike traditional intramuscular EMG from multiple muscles that distinguishes between fascicles that innervate different muscles but lacks spatial mapping within a muscle, HD-eEMG enables detailed characterization of motor unit activity and spatial activation patterns within a muscle (33–37). Our study shows that HD-eEMG combined with intrafascicular stimulation reveals spatially distinct and consistent muscle activation patterns, reflecting the selective recruitment of motor units based on electrode placement. This spatio-temporal information of M-wave responses to electrical stimulation allows the inference of activated fiber subpopulations and demonstrates reliable, repeatable results across experiments.

### 4.2 Selective activation of nerve fiber subpopulations can be achieved by altering electrode placement

In this study, we evaluated intrafascicular selectivity when using LIFEs and demonstrated that different subpopulations of nerve fibers within the same fascicle can be activated by altering electrode placement. Our results indicated that the muscle activation pattern varied depending on the electrode used for stimulation, despite all electrodes operating at comparable threshold levels (defined as the minimum current required to elicit a motor response). These variations in activation patterns can be attributed to the recruitment of different motor units (MUs) (33,36,38) that produce distinct motor response shapes across the HD-eEMG grid (37). These HD-eEMG findings were consistent with the observed experimental behavior, where localized quasi-contractions were evident in specific muscle regions, and these regions shifted when stimulation was delivered through a different electrode. These differences in eEMG waveform morphology are influenced by multiple factors, including the number of active MUs, dispersion of innervation zones, MU location, and MU distribution (37,39). These variations were observed across all HD-eEMG channels and AUC values were calculated from the profile curves.

Conventional PNS approaches predominantly use cuff electrodes that wrap the targeted nerve and deliver an electrical charge across the entire nerve trunk (4,13,40,41). Although cuff electrodes are less invasive and exhibit long-term stability (9,40–42), their limited spatial selectivity is a critical disadvantage. Cuff-based stimulation approaches often recruit multiple fiber subpopulations by stimulating a broad cross-section of the nerve, leading to unintended side effects (43). For example, PNS using cuff electrodes to restore sensory feedback in users of upper-limb prosthetics has been associated with unintended muscle activation (44), whereas using cuffs for vagus nerve stimulation can inadvertently activate the laryngeal muscles and reduce therapeutic tolerability (4,45,46).

In contrast, the geometric properties of LIFEs enable placement at various cross-sectional and longitudinal positions within a fascicle, thereby reducing the distance between the active site of the electrode and targeted nerve fibers (11,17). Fiber recruitment is governed by biophysical characteristics where large myelinated fibers exhibit lower activation thresholds than smaller, unmyelinated fibers, making electrode placement critical. LIFEs can be positioned in proximity to sensory, motor, or autonomic fibers, each characterized by distinct thresholds, conduction velocities, and functional roles (i.e., thermoreceptors, somatic motor neurons, visceral afferents, and nociceptors) (47,48). These spatial relationships enable intrafascicular stimulation to achieve higher selectivity than extraneural approaches, although the potential risk of activating non-targeted fibers owing to imprecise placement is heightened.

Furthermore, when the stimulation amplitude was increased in our experiments, motor responses exhibited a graded increase, reflecting the recruitment of additional subpopulations as the current spread within the fascicle. These findings are consistent with the well-known principles of nerve stimulation: increasing the stimulation amplitude, and thereby the total charge delivered, enables recruitment of nerve fibers with higher thresholds and expands the recruitment of fibers positioned farther from the electrode.

During this study, stimulation pulse trains were delivered to the gastrocnemius lateralis at tetanic frequencies low enough to produce a clear and reproducible muscle response while remaining below the threshold required to elicit maximal sustained contraction. This paradigm enabled the accurate assessment of muscle patterns and minimized fatigue by preventing complete contraction-relaxation cycles, thereby allowing consistent evaluation of M-wave responses. Although intrafascicular selectivity was assessed using muscle activity (HD-eEMG), these findings may be translatable to sensory fiber activation. Prior research has demonstrated that intraneural electrodes for the PNS can evoke precise and proportional sensory feedback in humans. For example, individuals with upper-limb amputations who have received intraneural implants have reported restoration of distinct tactile percepts and an enhanced ability to discriminate objects (7,18,44,49,50). The underlying biophysical principle remains unchanged: intrafascicular electrodes decrease the distance to the targeted axons, resulting in lower activation thresholds and limiting the spread of current. Therefore, the selectivity observed in motor activation is likely to be observed in sensory fibers. Hence, reducing the side effects of PNS-based therapies remains a significant challenge in the field of bioelectronic medicine. Enhancing the selectivity of the PNS paradigms is a promising approach for mitigating these undesirable effects.

Enhanced selectivity has the potential to improve the precision of the existing neurostimulation technologies. For instance, these results could inform PNS-based approaches for restoring sensory feedback in individuals with amputations by reducing off-target effects and improving and refining sensory inputs. Furthermore, improving selectivity may broaden the clinical and technological scope of PNS-based approaches, enabling the exploration of applications constrained by limited selectivity.

### 4.3 Electric field steering can be used for activation of additional fiber subpopulations

In our experiments, bipolar stimulation produced higher RMS values for motor responses than monopolar stimulation, indicating a stronger and more synchronized fiber activation. Although previous studies have suggested that bipolar stimulation decreases activation as electrode spacing increases (51), the spacing between multiple LIFEs in the sciatic nerve may have been insufficient to elicit this effect. Instead, these electrodes may act synergistically to generate a more concentrated electric field, resulting in an enhanced activation. The use of LIFEs in a bipolar configuration revealed principles comparable to those observed for extraneural electrodes. Multiple LIFEs implanted within the same fascicle used in a bipolar configuration provided access to fiber subpopulations that neither electrode could independently reach.

Electric field steering is an additional strategy for selective activation of nerve fiber subpopulations. By employing multiple stimulation sites and adjusting the polarity and amplitude, the spatial distribution of the current within the nerve can be directed toward specific fiber groups (21,23,52). Previous studies have used multi-contact cuff electrodes to enhance selectivity by shaping the electric field to target or avoid specific nerve regions. For example, Dali et al. (22) evaluated various multi-contact cuff configurations, including tripolar, longitudinal, transverse, and steering current rings, and demonstrated that altering these could steer the current to activate specific muscles, with each configuration exhibiting distinct effects on selectivity, efficiency, and charge propagation. Computational modeling studies have similarly suggested that electric field steering could optimize vagus nerve stimulation to modulate heart rate while reducing side effects compared to standard ring cuff configurations (53).

Our bipolar configuration findings demonstrate that implanting multiple LIFEs within the same fascicle could enable additional stimulation strategies that cannot be achieved with a single electrode. By alternating and evaluating different electrode combinations, it is possible to refine the stimulation paradigms tailored to specific recruitment patterns. Ultimately, this approach may enhance selectivity by steering activation across multiple electrodes, thus providing a more precise and adaptable method for targeting distinct fiber subpopulations.

## 5. Limitations

Although LIFEs have been successfully implanted in humans for clinical applications, such as restoration of sensory feedback in individuals with amputations, there are currently no surgical techniques that enable precise implantation of intraneural electrodes to target a pre-specified fiber population. Currently, this limitation extends to general applications but was also present in our studies. Implantation of intrafascicular electrodes typically relies on morphological data from prior studies (54,55) without real-time confirmation of electrode positioning. As demonstrated in this study, electrode placement is critical for achieving intrafascicular selectivity; even minor positional differences can substantially alter the subset of activated fibers. Therefore, the development of surgical techniques or guidance systems that allow high-precision electrode placement can significantly reduce the off-target effects.

Furthermore, although LIFEs have been used in clinical trials, they have not yet received FDA approval as a therapeutic neural interface. Consequently, there are a limited number of research groups working on improving implantation techniques or optimizing stimulation paradigms that use LIFEs.

## 6. Conclusions

Enhancing the selectivity of peripheral nerve stimulation to minimize undesirable side effects continues to pose a significant challenge to bioelectronic medicine. Using multiple longitudinal intrafascicular electrodes implanted within the sciatic nerve, we investigated the effects of intrafascicular electrode placement and field steering on stimulation selectivity. We demonstrate that HD-eEMG is a robust method for evaluating intrafascicular selectivity. Our results indicate that both approaches are capable of selectively activating discrete subpopulations of nerve fibers within a fascicle, thereby improving stimulation specificity and offering the potential to reduce off-target activation in bioelectronic medical therapies.

## 7. Declarations

### Ethical Approval and Consent to Participate

The Institutional Animal Care and Use Committee of the University of Arkansas approved the study procedures and protocols (Protocol AUP22014).

### Data Availability

The data supporting the findings of this study are openly available at: https://doi.org/10.5061/dryad.rbnzs7hth

### Authors’ contributions

A.O.S., A.H., A.K.T. and L.R. contributed to the data collection. A.O.S., L.R, J.J.A. and L.M. contributed to data analysis. A.O.S., S.R., L.M., and J.M.A. contributed to statistical analyses. F.K., Y.B., and L.R. contributed to the design, and construction of the stimulation hardware and the development and programming of associated software algorithms. A.O.S. wrote the original draft of the manuscript. J.J.A. and R.J. reviewed all aspects of the research including the writing. R.J., J.J.A., S.R., Y.B., and F.K. conceptualized the project and secured funding. All authors reviewed, read, and approved the final manuscript.

## Acknowledgments

Not applicable

